# Bacterial biodiversity drives the evolution of CRISPR-based phage resistance in *Pseudomonas aeruginosa*

**DOI:** 10.1101/586115

**Authors:** Ellinor O Alseth, Elizabeth Pursey, Adela M Luján, Isobel McLeod, Clare Rollie, Edze R Westra

## Abstract

Approximately half of all bacterial species encode CRISPR-Cas adaptive immune systems^1^, which provide immunological memory by inserting short DNA sequences from phage and other parasitic DNA elements into CRISPR loci on the host genome^2^. Whereas CRISPR loci evolve rapidly in natural environments^3^, bacterial species typically evolve phage resistance by the mutation or loss of phage receptors under laboratory conditions^4,5^. Here, we report how this discrepancy may in part be explained by differences in the biotic complexity of *in vitro* and natural environments^6,7^. Specifically, using the opportunistic pathogen *Pseudomonas aeruginosa* and its phage DMS3*vir*, we show that coexistence with other human pathogens amplifies the fitness trade-offs associated with phage receptor mutation, and therefore tips the balance in favour of CRISPR-based resistance evolution. We also demonstrate that this has important knock-on effects for *P. aeruginosa* virulence, which became attenuated only if the bacteria evolved surface-based resistance. Our data reveal that the biotic complexity of microbial communities in natural environments is an important driver of the evolution of CRISPR-Cas adaptive immunity, with key implications for bacterial fitness and virulence.

---

*Pseudomonas aeruginosa* is a widespread opportunistic human pathogen that thrives in a range of different environments, including hospitals, where it is a common source of nosocomial infections. In particular, it frequently colonises the lungs of cystic fibrosis patients, in whom it is the leading cause of morbidity and mortality^8^. In part fuelled by a renewed interest in the therapeutic use of bacteriophages as antimicrobials (phage therapy)^9,10^, many studies have examined if and how *P. aeruginosa* evolves resistance to phage (reviewed in ref. 11). The clinical isolate *P. aeruginosa* strain PA14 has been reported to predominantly evolve resistance against its phage DMS3*vir* by the modification or complete loss of the phage surface receptor^12^ when grown in nutrient-rich medium^4^ despite carrying an active CRISPR-Cas adaptive immune system (<u>C</u>lustered <u>R</u>egularly Interspaced <u>S</u>hort <u>P</u>alindromic <u>R</u>epeats; <u>C</u>RISPR-<u>as</u>sociated). Conversely, under nutrient-limited conditions, the same strain relies on CRISPR-Cas to acquire phage resistance^4^. While these observations suggest abiotic factors are critical determinants of the evolution of phage resistance strategies, the role of biotic factors has remained unclear, even though *P. aeruginosa* commonly co-exists with a range of other bacterial species in both natural and clinical settings^13,14^.

To explore how microbial biodiversity impacts the evolution of phage resistance, we co-cultured *P. aeruginosa* PA14 with three other clinically relevant opportunistic pathogens that are known to co-infect with *P. aeruginosa*, namely *Staphylococcus aureus, Burkholderia cenocepacia*, and *Acinetobacter baumannii*^14–16^, none of which can be infected by or interact with phage DMS3*vir* (Extended Data Fig. 1, Linear model: Effect of *P. aeruginosa* on phage titre over time; t = −3.37, p < 0.001; *S. aureus*; t = 1.63, p = 0.11; *A. baumannii*; t = 1.20, p = 0.23; *B. cenocepacia*; t = −0.27, p = 0.79; Overall model fit; F_9,235_ = 4.33, adjusted R^2^ = 0.11, p < 0.001). We applied a “mark-recapture” approach using a *P. aeruginosa* PA14 mutant carrying streptomycin resistance in order to monitor the bacterial population dynamics and phage resistance evolution in the focal subpopulation at 3 days post infection (d.p.i.). This revealed that in nutrient-rich broth (Lysogeny Broth), PA14 evolved significantly higher levels of CRISPR-based resistance following infection with 10^6^ plaque forming units (p.f.u.) of phage DMS3*vir* when co-cultured with other bacterial species compared to when grown in isolation or co-cultured with an isogenic surface mutant (Fig. 1). Additionally, we found that these effects were dependent on the identity of the species that were present in the mixed culture, with the strongest effects being observed in the presence of *A. baumannii* or a mix of the three bacterial species, and an absence of any effect when PA14 was co-cultured with an isogenic surface mutant that lacked the phage receptor (Fig. 1, Deviance test: Relationship between community composition and CRISPR; Residual deviance(30, n = 36) = 1.81, p < 0.001; Tukey contrasts: Monoculture v Mixed; z = −5.99, p < 0.001; Monoculture v *A. baumannii*; z = −4.33, p < 0.001; Monoculture v *B. cenocepacia*; z = −3.76, p < 0.01; Monoculture v *S. aureus*; z = −2.38, p = 0.14; Monoculture v surface mutant; z = 2.26, p = 0.19). To explore the clinical relevance of this observation, we next co-cultured *P. aeruginosa* with *A. baumannii* in artificial sputum medium (ASM), which mimics the abiotic environment of sputum from cystic fibrosis patients^17^. This analysis revealed a similar effect of *A. baumannii* on the evolution of CRISPR-based resistance in both LB and ASM (Extended Data Fig. 2, Deviance test: Relationship between community composition and CRISPR; Residual deviance(20, n = 22) = 2.33, p < 0.001; Relationship between growth medium and CRISPR; Residual deviance(19, n = 22) = 2.30, p = 0.59). To further explore the generality of our finding, we next manipulated the microbial community composition by varying the proportion of *P. aeruginosa* versus the other pathogens. This revealed that increased CRISPR-based resistance evolution occurred across a wide range of microbial community compositions, with a maximum effect size when *P. aeruginosa* made up 50% of the initial bacterial mixture (Extended Data Fig. 3). An exception to this trend was when the *P. aeruginosa* subpopulation made up only 1% of the total community; in this case sensitive bacteria persisted due to the reduced size of the phage epidemic and hence relaxed selection for resistance (Extended Data Fig. 3). Collectively, these data suggest that greater levels of interspecific competition contribute to the evolution of CRISPR-based resistance.

**Figure 1:**
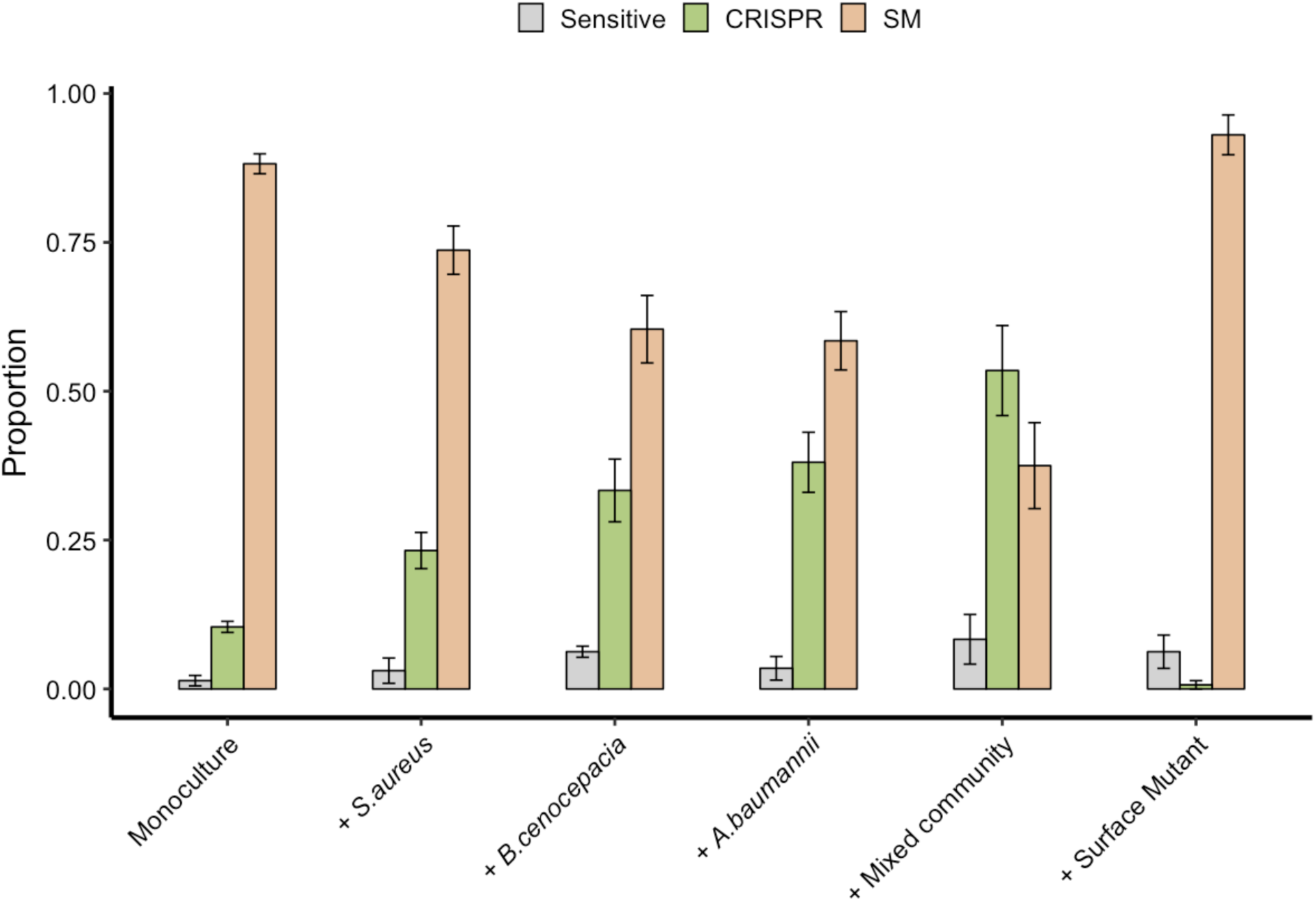
Biodiversity affects the evolution of phage resistance. Proportion of *P. aeruginosa* that acquired surface modification (SM) or CRISPR-based resistance, or remained sensitive at 3 days post infection with phage DMS3*vir* (n = 6 for all treatments). *P. aeruginosa* was grown in monoculture and in polycultures with other bacterial species individually or as a mixture, or with an isogenic surface mutant. Error bars correspond to ± one standard error.

Given that cell surface molecules likely play a part in interspecific competition^18^, we hypothesised that the fitness cost of surface-based resistance may be amplified in the presence of other bacterial species, resulting in stronger selection for bacteria with CRISPR-based resistance. To test this, we competed the two phage resistant phenotypes (i.e. CRISPR-resistant and surface mutant clones) in the presence or absence of the microbial community, and across a range of phage titres. In the absence of the microbial community and phage, CRISPR-resistant bacteria showed a small fitness advantage over bacteria with surface-based resistance, but this advantage disappeared when phage was added and as titres increased (Fig. 2a, and ref. 7). In the presence of the biodiverse microbial community however, the relative fitness of bacteria with CRISPR-based resistance was consistently higher, demonstrating that mutation of the Type IV pilus is more costly when bacteria compete with other bacterial species (Fig. 2a, Linear model: Effect of community absence; t = −5.54, p < 0.001; Effect of increasing phage titre; t = −2.41, p < 0.05; Overall model fit; Adjusted R^2^ = 0.41, F_4,139_ = 25.48, p < 0.001). This increased fitness trade-off associated with surface-based resistance was also observed when the CRISPR- and surface-resistant phenotypes competed in the presence of only a single additional species (Fig. 2b, ANOVA with Tukey contrasts: Overall difference in fitness; F_4,2_ = 8.151 p < 0.001; Monoculture v Mixed; p < 0.05; Monoculture v *A. baumannii*; p < 0.05; Monoculture v *B. cenocepacia*; p < 0.05), with the exception of *S. aureus* (Fig. 2d. Monoculture v *S. aureus*; p = 0.80), concordant with this species inducing the lowest levels of CRISPR (Fig. 1). These fitness trade-offs therefore explain why *P. aeruginosa* evolved greater levels of CRISPR-based resistance in the presence of the other pathogens, and why this varied depending on the species (Fig. 1).

**Figure 2:**
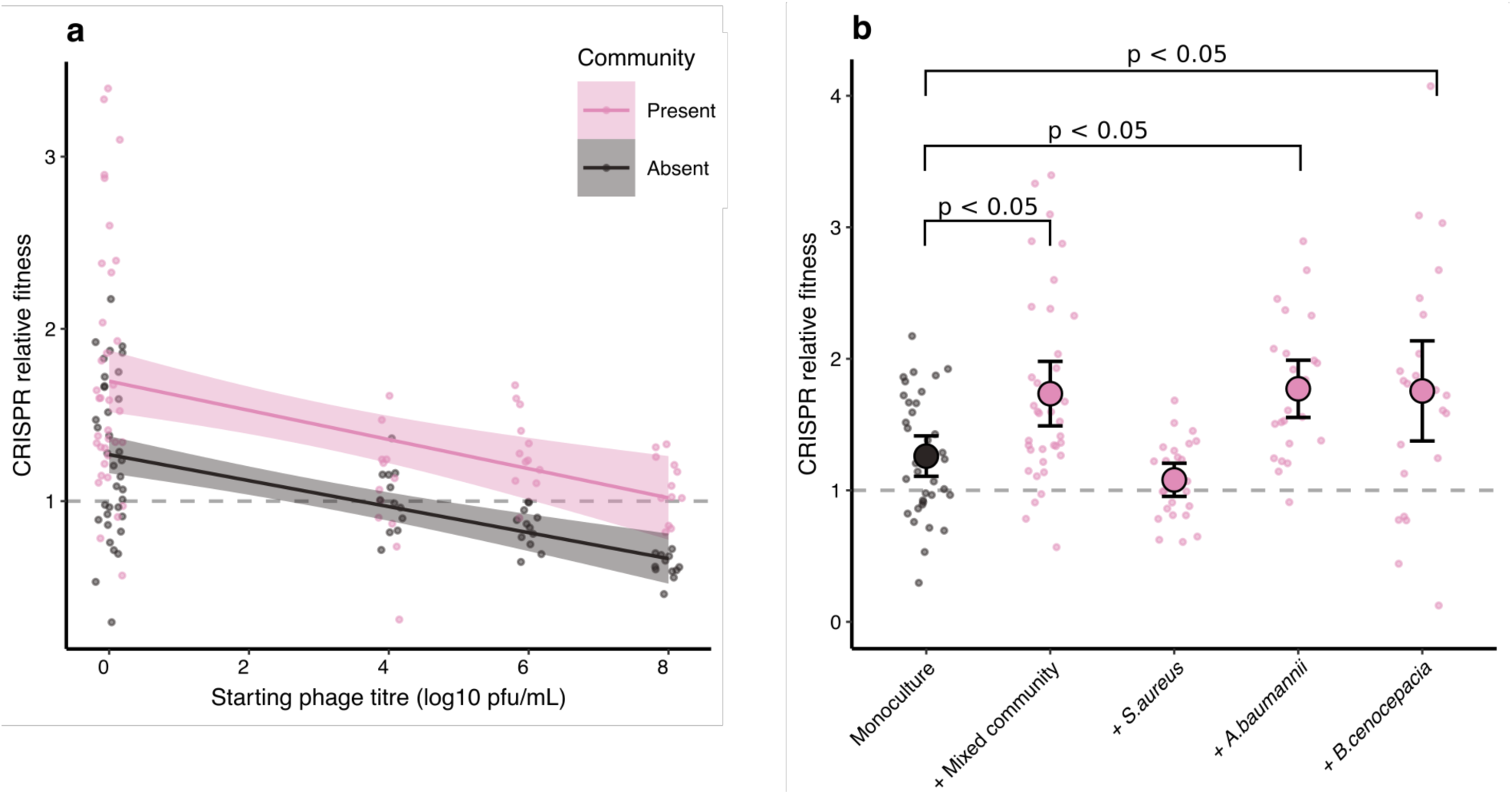
Biodiversity mitigates fitness costs associated with CRISPR-based resistance. Relative fitness of a *P. aeruginosa* clone with CRISPR-based resistance after competing for one day against a surface modification clone at (**a**) varying levels of phage DMS3*vir* in the presence or absence of a mixed microbial community, or (**b**) in absence of phage, but in the presence of a mixed community or its individual bacterial species. Error bars correspond to 95% confidence intervals.

Evolution of phage resistance by bacterial pathogens is often associated with virulence trade-offs when surface structures are modified^19–21^, whereas similar trade-offs have not yet been reported in the literature for CRISPR-based resistance. We therefore hypothesised that the community context in which phage resistance evolves may have important knock-on effects for *P. aeruginosa* virulence. To test this, we used a *Galleria mellonella* larvae infection model, which is commonly used to evaluate the virulence of human pathogens^22,23^. We compared *in vivo* virulence of *P. aeruginosa* clones that evolved phage resistance in different community contexts by injecting larvae with a mixture of clones that had evolved phage-resistance in either the presence or absence of the mixed bacterial community (Extended Data. Fig. 2). Taking time to death as a proxy for virulence, we found that the evolution of phage resistance in the presence of a microbial community was associated with greater levels of *P. aeruginosa* virulence compared to when phage-resistance evolved in monoculture, and similar to that of the ancestral PA14 strain (Fig. 3a, Cox proportional hazards model with Tukey contrasts: Community present v absent; z = 5.85, p < 0.001; ancestral PA14 v community absent; z = 4.42, p < 0.001; ancestral PA14 v community present; z = −1.30, p = 0.38. Overall model fit; LRT_3_ = 51.03, n = 376, p < 0.001). These data, in combination with the fact that the Type IV pilus is a well-known virulence factor^19,24^, are consistent with the idea that the mechanism by which bacteria evolve phage resistance has important implications for bacterial virulence. To more directly test this, we next infected larvae with each individual *P. aeruginosa* clone for which we had previously determined the mechanism underlying evolved phage resistance (Extended Data Fig. 2), again using time to death as a measure of virulence. This showed that bacterial clones with surface-based resistance both had drastically reduced swarming motility (consistent with mutations in the Type IV pilus^19^) (Fig. 3b, ANOVA with Tukey contrasts: Overall effect; F_2,977_ = 472.5, p < 0.001; Sensitive v CRISPR; p = 0.88; CRISPR v Surface mutant; p < 0.001) and impaired virulence (Fig. 3c, Cox proportional hazards model with Tukey contrasts: Surface mutant v CRISPR; z = −2.37, p < 0.05; Sensitive v CRISPR; z = 2.10, p = 0.10; Surface mutant v Sensitive; z = −4.23, p < 0.001. Overall model fit; LRT_3_ = 48.66, n = 981, p < 0.001).

**Figure 3:**
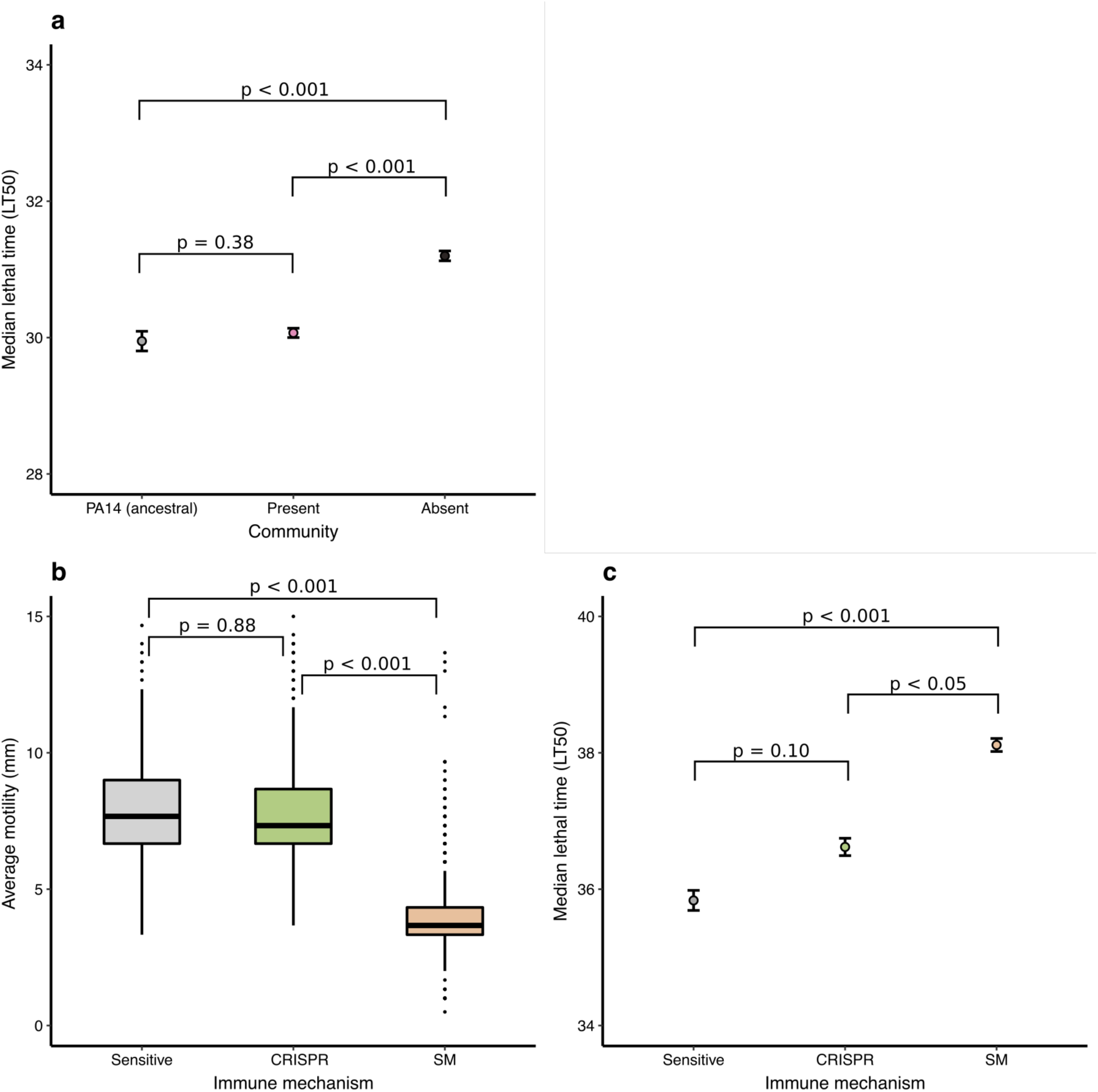
Evolution of phage resistance affects *in vivo* virulence. Median lethal time (LT50 = the timepoint when half of the infected larvae were dead) for (**a**) PA14 clones that evolved phage resistance either in the presence or absence or a mixed microbial community. Type of evolved phage resistance drastically impacted (**b**) motility, and (**c**) *in vivo* virulence. Error bars correspond to ± one standard error.

Collectively, our data show that the evolutionary outcome of bacteria-phage interactions can be fundamentally altered by the microbial community context. While traditionally studied in isolation, these interactions are usually embedded in complex biotic networks of multiple species, and it is becoming increasingly clear that this can have key implications for the evolutionary epidemiology of infectious disease, including the evolution of pathogen virulence and host range^25–29^. The work presented here reveals that the community context can also shape the evolution of different host resistance strategies. Specifically, we find that the interspecific interactions between four bacterial species in a synthetic microbial community can have a large impact on the evolution of phage resistance mechanisms by amplifying the constitutive fitness cost of surface-based resistance^4^. The finding that biotic complexity matters complements previous work on the effect of abiotic variables on phage resistance evolution. This previous work showed that a greater force of infection resulted in a higher induced fitness cost of CRISPR-based resistance, hence selecting for surface-based resistance in the presence of higher phage titres^4^. Variation in the force of infection did not however seem to play a role in the effects described in this study, since even though phage epidemic sizes varied depending on the microbial community composition (Extended Data Fig. 4, ANOVA: Overall effect of *P. aeruginosa* starting percentage on phage titre; F_6,105_ = 14.84, p < 0.001), they did not correlate with the levels of evolved CRISPR-based resistance (Extended Data Fig. 5). This therefore suggests that in this case the impact of biotic complexity on the evolution of CRISPR-based resistance is stronger than that of variation in phage abundance. While future work will be critical to further generalise the findings described here to other bacterial species and strains, we speculate that the way in which the microbial community composition drives the evolution of phage resistance strategies may be important in the context of phage therapy. Primarily, the absence of detectable trade-offs between CRISPR-based resistance and virulence, as opposed to when bacteria evolve surface-based resistance, suggests that evolution of CRISPR-based resistance can ultimately influence the severity of disease. Moreover, evolution of CRISPR-based resistance can drive more rapid phage extinction^30^, and may in a multi-phage environment result in altered patterns of cross-resistance evolution compared to surface-based resistance^31^. The identification of the drivers and consequences of CRISPR-resistance evolution might help to improve our ability to predict and manipulate the outcome of bacteria-phage interactions in both natural and clinical settings.

## Acknowledgements

The authors thank Prof. A. Buckling for critical reading of the manuscript, and Prof. JP Pirnay and D. de Vos for sharing clinical isolates of *S. aureus, A. baumannii*, and *B. cenocepacia*. This work was supported by grants from the ERC (ERC-STG-2016-714478 - EVOIMMECH) and the NERC (NE/M018350/1), which were awarded to E.R.W.

## Author Contributions

Conceptualisation: E.O.A. and E.R.W. Methodology: E.O.A., A.M.L., C.R. and E.R.W. Investigation: E.O.A., E.P., I.M. Formal analysis: E.O.A., E.P., I.M., C.R. and E.R.W. Writing – Original draft: E.O.A. Writing – Review and editing: E.O.A. and E.R.W. Funding acquisition: E.R.W.

## Materials and correspondence

All materials used in this study are available upon request to Edze Westra or Ellinor Alseth.

## Methods

All statistical analyses were done using R version 3.5.1. (R Core Team, 2018), and the Tidyverse package version 1.2.1. (Wickham, 2017). All *Galleria mellonella* mortality analyses were done using the Survival package version 2.38 (Therneau, 2015).

### Bacterial strains and viruses

We used a marked *P. aeruginosa* UCBPP-PA14 mutant carrying a streptomycin resistant gene inserted into the genome using pBAM1^32^ (referred to as the ancestral PA14 strain). The WT PA14 bacteriophage-insensitive mutant with 2 CRISPR spacers (BIM5), the surface mutant derived from the PA14 *csy3::LacZ* strain, and phage DMS3*vir* and DMS3*vir* +*acrF1*(carrying an anti-CRISPR gene) have all been previously described (refs. 4 and 29 and references therein). The bacteria used as the microbial community were *Staphylococcus aureus* strain 13 S44 S9, *Acinetobacter baumannii* clinical isolate FZ21 and *Burkholderia cenocepacia* J2315, were all isolated from patients at Queen Astrid Military Hospital, Brussels, Belgium.

### Absorption and infection assays

Phage infectivity against each of the bacterial species used in this study was assessed by spotting serial dilutions of virus DMS3*vir* on lawns of the individual community bacteria, followed by checking for any plaque formation after 24 hours of growth at 37°C. Adsorption assays (as shown in Extended Data Fig. 1) were performed by monitoring phage titres over time, for up to an hour (At 0, 2, 4, 6, 8, 10, 15 and 20 minutes post infection for PA14, and at 0, 5, 10, 20, 40 and 60 minutes post infection for the individual microbial community bacteria), after inoculating the individual bacteria in mid-log phage at approximately 2 × 10^8^ c.f.u. with phage DMS3*vir* at 2 × 10^6^ p.f.u. (final MOI = 0.001). Adsorption assays were carried out in LB medium, incubated at 37°C while shaking at 180 r.p.m. (three independent replicates per experiment).

### Coevolution experiments

The streptomycin resistant ancestral strain of *P. aeruginosa* was used for all coevolution experiments. The experiments (shown in Fig. 1, and Extended Data Figs. 2 and 3) were performed by inoculating 60μl of approximately 10^6^ colony-forming units (c.f.u.) of bacteria from overnight cultures into glass microcosms containing 6ml LB medium (Fig. 1 and Extended Data Fig. 3), or artificial sputum medium (for composition, see ref. 17) (Extended Data Fig. 2), followed by incubation at 37°C while shaking at 180 r.p.m. (n= 6 per treatment, expect for Extended Data Fig. 2 monoculture in LB treatment, where n = 4). The polyculture mixes either consisted of approximately equal amounts of all four bacterial species or mixes of *P. aeruginosa* with just one additional species where *P. aeruginosa* made up 25% of the total volume (60µl). For the experiment shown in Extended Data Fig. 3, *P. aeruginosa* was grown in the presence of the mixed microbial community, but at a range of different starting percentages based a total volume of 60µl. Before inoculation, phage DMS3*vir* was added at 10^6^ p.f.u. (Fig. 1), or at 10^4^ p.f.u. (Extended Data Fig. 2 and 3). Transfers of 1:100 into fresh broth were done daily for a total of three days. Additionally, phage titres were monitored daily by spotting chloroform-extracted phage dilutions on a lawn of *P. aeruginosa csy3::LacZ*. Downstream analysis to determine phage resistance was done using cross-streak assays and PCR as described in ref. 4.

### Competition experiments

For both competition experiments shown in Fig. 2, the BIM5 clone was competed against the *csy3::LacZ* surface mutant. Bacteria were grown for 24 hours in glass microcosms containing 6ml LB medium, in a shaking incubator at 180 r.p.m. and at 37^°^C. For the experiment shown in Fig. 2a, the two phenotypes were competed in the presence or absence of the mixed microbial community, either without the addition of phage (n = 36), or infected with phage DMS3*vir* at 10^4^, 10^6^, and 10^8^ p.f.u. (n = 12 per treatment). For the experiment shown in Fig. 2b, the two phage resistant phenotypes were again competed either in the presence or absence of the mixed microbial community, with the addition of the individual bacterial species with the *P. aeruginosa* phenotypes always making up 25% of the total volume of 60µl (n = 24 per treatment). Samples were taken at 0 and 24 hours post infection., and the cells were serial diluted in M9 salts and plated on cetrimide agar (Sigma) supplemented with ca. 50µg ml-1 X-gal (to select for *P. aeruginosa*, while also differentiating between the BIM5 CRISPR clones (white) and CRISPR-KO surface mutant (blue)). Relative fitness was calculated as described in refs. 4 and 29.

### *Galleria mellonella* infection experiments

All virulence assays were done by injecting 10µl of diluted sample into the rear proleg of individual *Galleria mellonella* larvae using a sterile syringe as described in ref. 23. All larvae were checked for mortality and melanisation before injection. For the experiment shown in Fig. 3a, all evolved clones from the 25% (community present) and 100% (community absent) (Extended Data Fig. 3) were pooled together by replica (n = 6 per treatment) in 6mL of LB medium, with ten larvae infected per replica in three independent repeats (total no. of larvae = 376). To assess virulence of all evolved clones (Fig. 3c), infections were done independently using all the individual PA14 clones from the experiment shown in Extended Data Fig. 3, n = 981. Throughout the experiment, larvae were stored in 12-well plates, with one larva per well. Prior to injection, inoculums were prepared by transferring 5µl of all clones from the coevolution experiment to new 96-well plates containing 200µl LB medium, before being incubated for 24h at 37°C on an orbital shaker (180 r.p.m). For all experiments, after overnight growth at 37°C rotating at 180 r.p.m., the bacteria were diluted by adding 20µl to 180µl of M9 salts. Cell density was then assayed by measuring OD_600_ absorbance, with 0.1OD being ∼1 × 10^8^ cfu/ml, before being further diluted down to approximately 10^4^ cfu/ml, which was subsequently used for injection. Following infection, larvae were incubated at 28°C, with mortality monitored hourly for up to 48 hours. For both experiments, a control where larvae were injected with just M9 salts was included.

### Motility assays

Swarming motility of all evolved bacterial clones from the experiment shown in Extended Data Fig. 3 (n = 980) was assayed by using a 96-well microplate pin replicator to stamp the individual clones on 1% agar before overnight growth at 37°C. The diameters of the individual clones were then taken as a measure of motility (three replicas per clone).

---

**Extended Data Figure 1.**
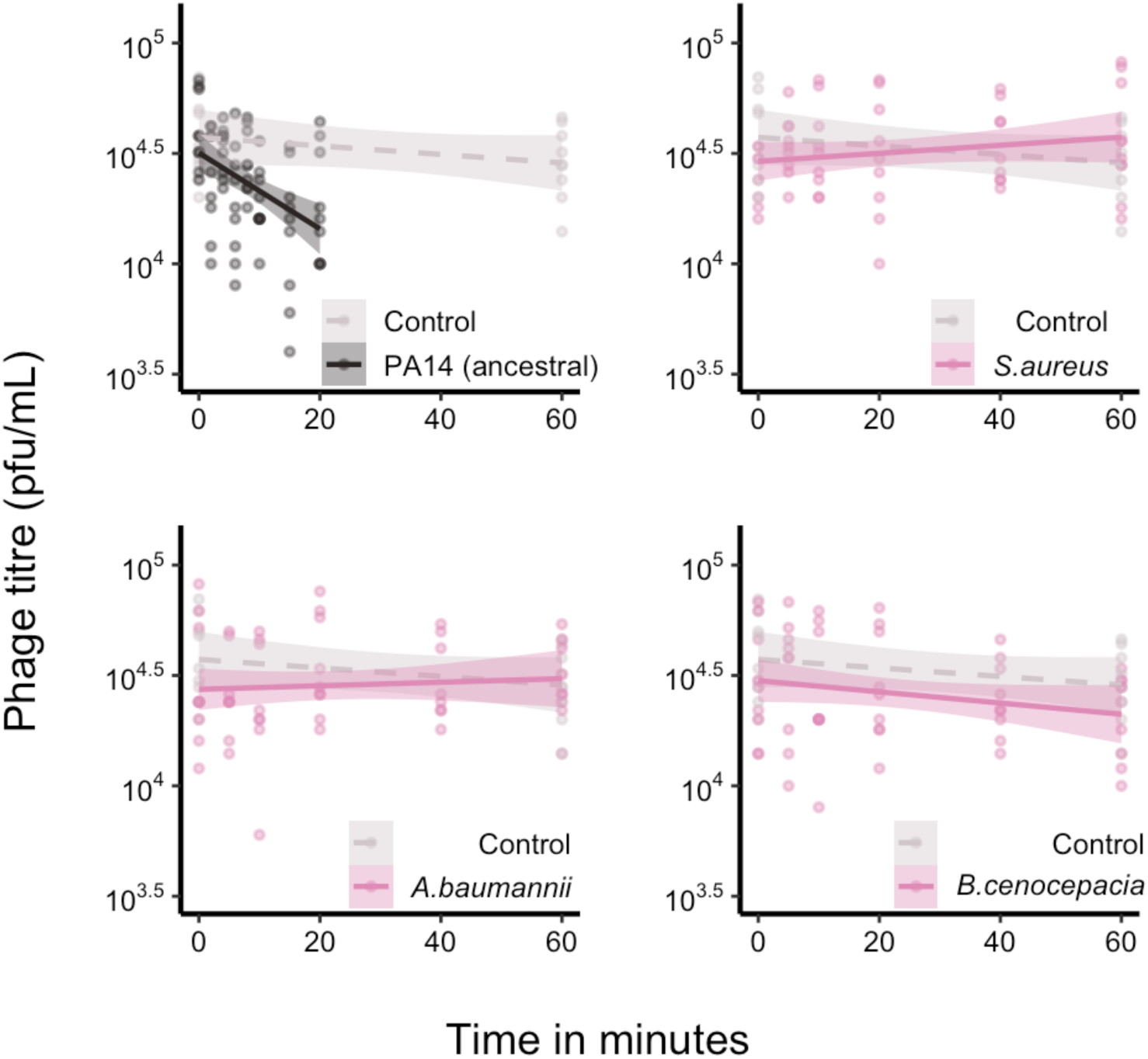
Only *P. aeruginosa* adsorbs phage DMS3*vir.* Phage levels, given in plaque-forming units per millilitre, in minutes post infection of *P. aeruginosa* PA14 and three other bacterial species. Controls were carried out in the absence of bacteria. Shaded areas correspond to 95% confidence intervals.

**Extended Data Figure 2.**
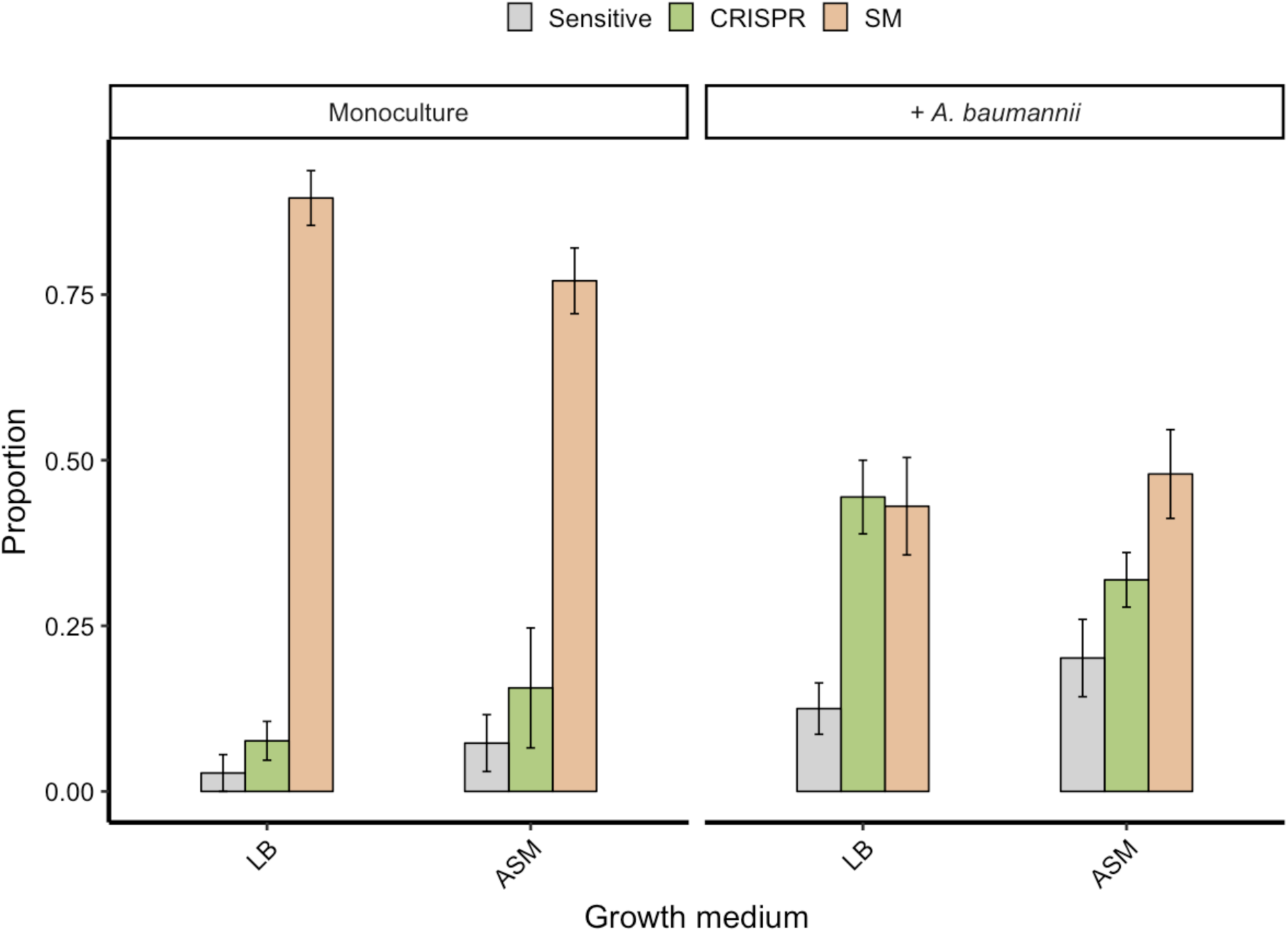
Enhanced CRISPR resistance evolution in artificial sputum medium. Proportion of *P. aeruginosa* that acquired surface modification (SM) or CRISPR-based immunity (or remained sensitive) at 3 days post infection with phage DMS3*vir* when grown in either LB or artificial sputum medium (ASM), and in the absence or presence of *A. baumannii*. Error bars correspond to ± one standard error.

**Extended Data Figure 3.**
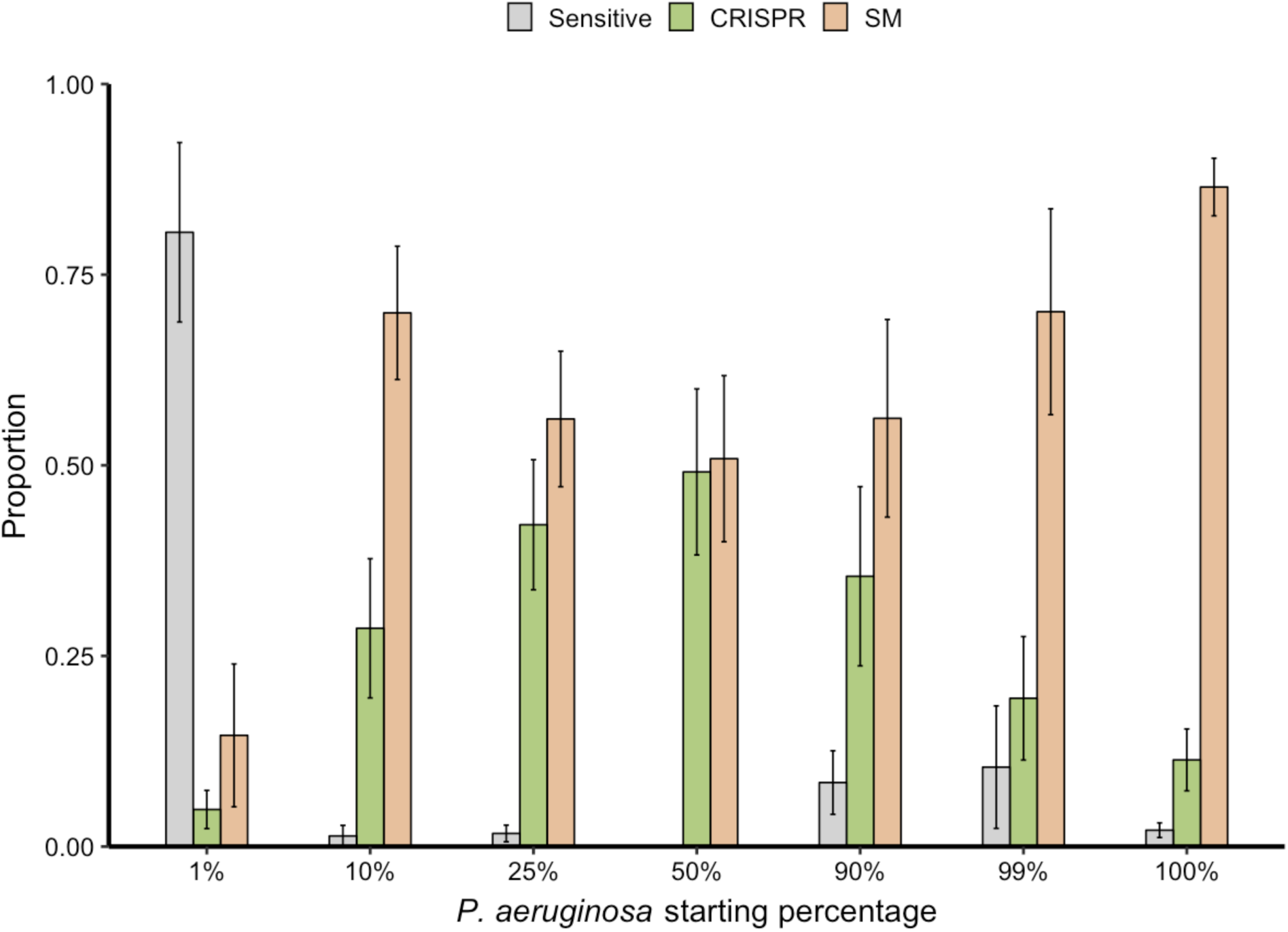
Increased CRISPR-based resistance evolution across a range of microbial community compositions. Proportion of *P. aeruginosa* that acquired surface modification (SM) or CRISPR-based immunity (or remained sensitive) at 3 days post infection with phage DMS3*vir* when grown either in monoculture (100%), or in polyculture mixtures consisting of the mixed microbial community but with varying starting percentages of *P. aeruginosa* based on volume. Error bars correspond to ± one standard error. Deviance test: Relationship between CRISPR and *P. aeruginosa* starting percentage; Residual deviance(35, n = 42) = 8.24, p < 0.001; 1%; z = −3.38, p < 0.01; 10%; z = 2.12, p < 0.05; 25%; z = 2.77, p < 0.01; 50%; z = 3.07, p < 0.01; 90%; z = 2.46, p < 0.05; 99%; z = 1.55, p = 0.13; 100%; z = 0.87, p = 0.39.

**Extended Data Figure 4.**
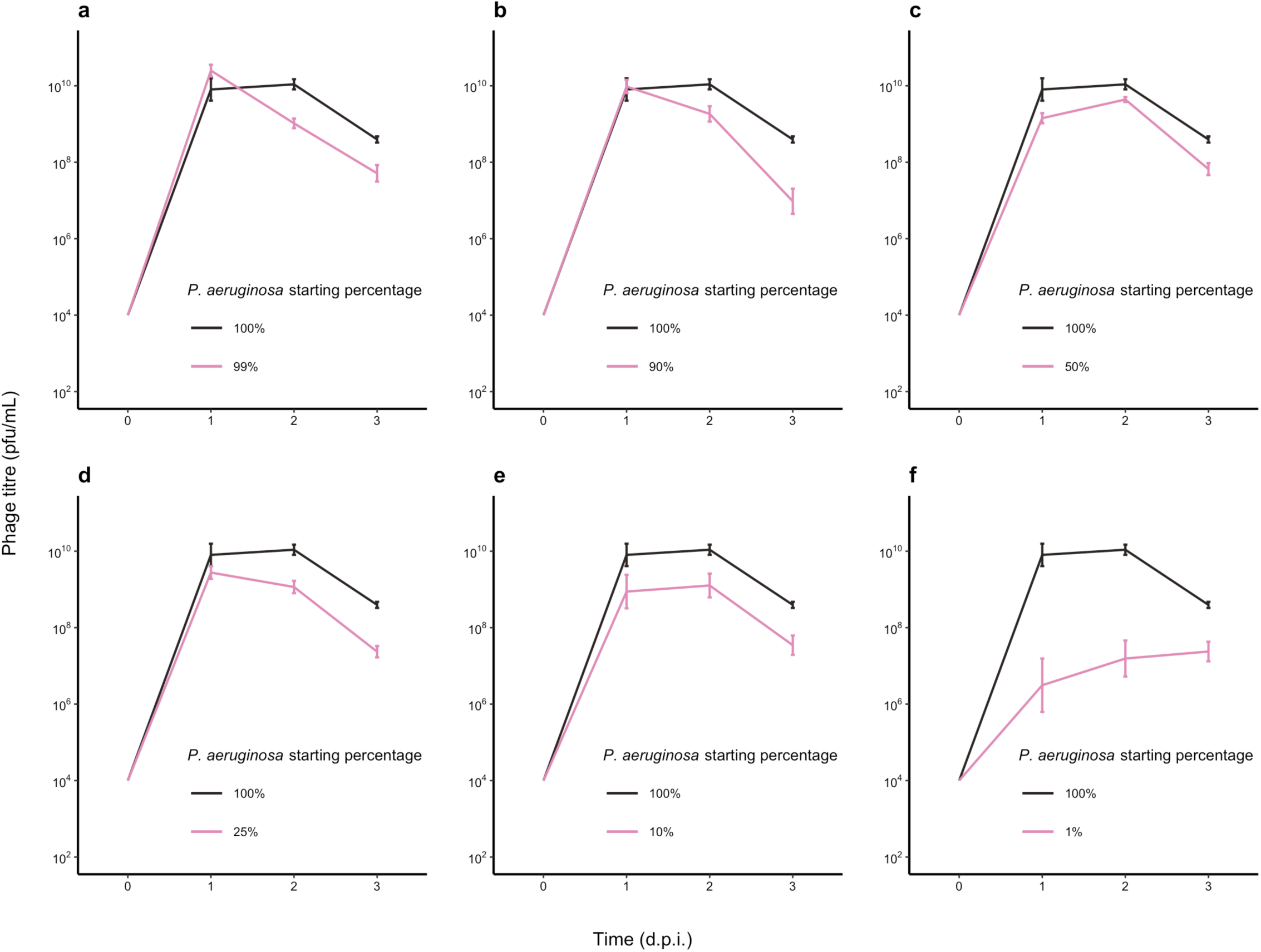
Microbial community composition impacts phage epidemic size. The DMS3*vir* phage titres (in plaque-forming units per millilitre) over time up to 3 days post infection of *P. aeruginosa* grown either in monoculture (100%), or in polyculture mixtures as shown in Extended Data Fig. 3. Error bars correspond to ± one standard error.

**Extended Data Figure 5.**
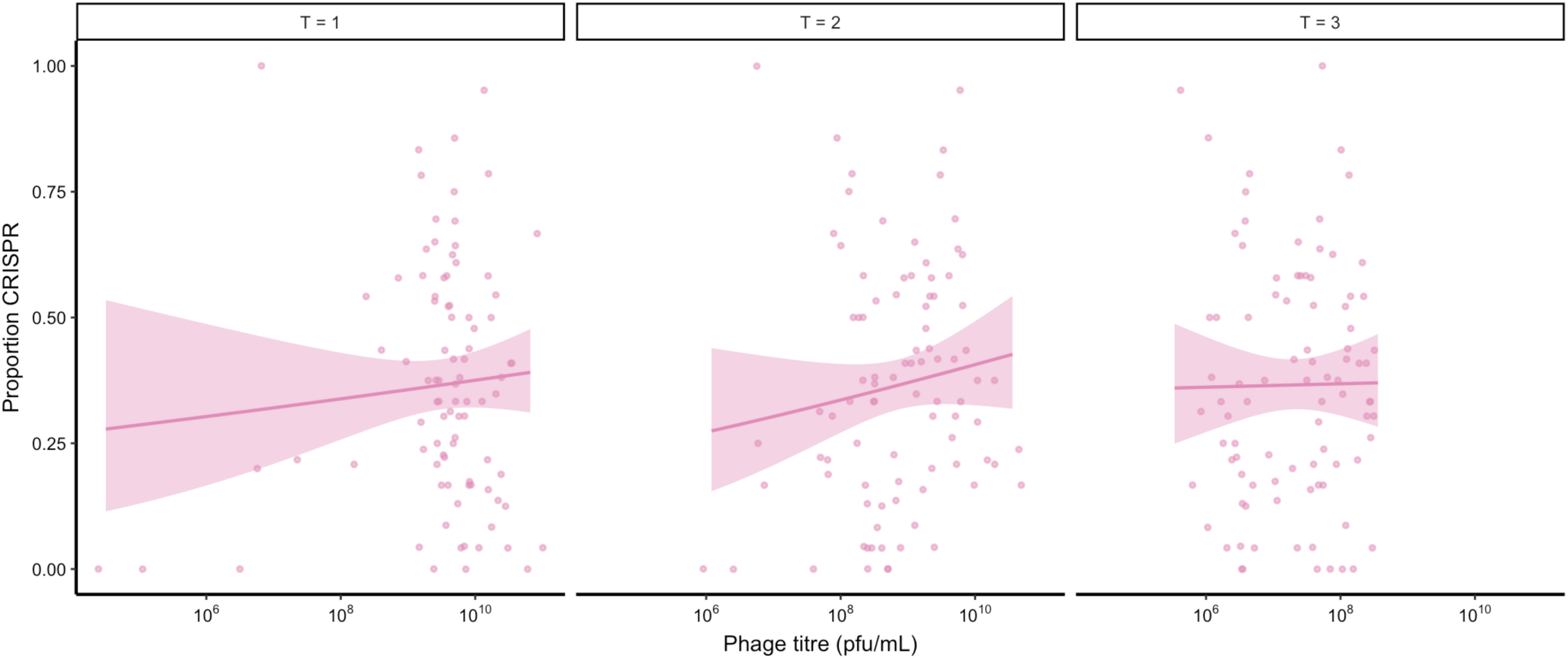
No correlation between phage epidemic size and evolution of CRISPR resistance. The correlation between the proportion of evolved CRISPR-based resistance and the phage epidemic sizes (in plaque-forming units per millilitre) using data taken from experiments shown in Fig. 1, Extended Data Fig. 2 and Extended Data Fig. 3. Correlations are separated by day, as phage titre was measured daily. Shaded areas correspond to 95% confidence intervals. Pearson’s correlations between phage titres (at each day post infection) and levels of CRISPR-based resistance.: T = 1; t_88_ = 0.75, p = 0.50, R^2^ = 0.08; T = 2; t_88_ = 1.21, p = 0.23, R^2^ = 0.13; T = 3; t_88_ = 0.11, p = 0.92, R^2^ = 0.01.

